# MethylCurate: Tool for Dataset Curation and Epigenetic Aging Clock Evaluation

**DOI:** 10.64898/2026.05.11.723515

**Authors:** Travyse A Edwards, Qi Long, Li Shen

## Abstract

**Summary:** DNA methylation datasets from public repositories such as NCBI Gene Expression Omnibus are central to the development and evaluation of epigenetic aging clocks, yet existing resources and tools do not fully resolve the bottlenecks of dataset retrieval and metadata harmonization. Current benchmarking frameworks often rely on static curated collections, support only a subset of available Gene Expression Omnibus studies, focus on specific tissues, or require substantial manual intervention when metadata fields and supplementary files are inconsistently structured across studies. We developed MethylCurate, an agentic AI framework that addresses these limitations by automating the retrieval of DNA methylation datasets from the Gene Expression Omnibus, harmonizing heterogeneous metadata, mapping datasets to a unified format, and enabling scalable evaluation of epigenetic aging clocks through an integrated, dialogue-driven workflow.

**Availability and Implementation:** MethylCurate is implemented in Python and combines deterministic modules for Gene Expression Omnibus dataset retrieval, quality control, and clock evaluation with large language model–assisted agents for metadata extraction, metadata harmonization, and DNA methylation data parsing. Source code, documentation, and example workflows are available at: https://github.com/Travyse/methylcurate

**Supplementary Information:** Supplementary data are available at Bioinformatics online.

**Graphical Abstract:** MethylCurate is an agentic-AI framework that converts user-specified NCBI Gene Expression Omnibus DNA methylation datasets into standardized metadata, beta matrices, artifacts, logs, and aging clock benchmarking outputs through automated retrieval, quality control, metadata extraction, harmonization, and evaluation workflows. Figure generated with Biorender.

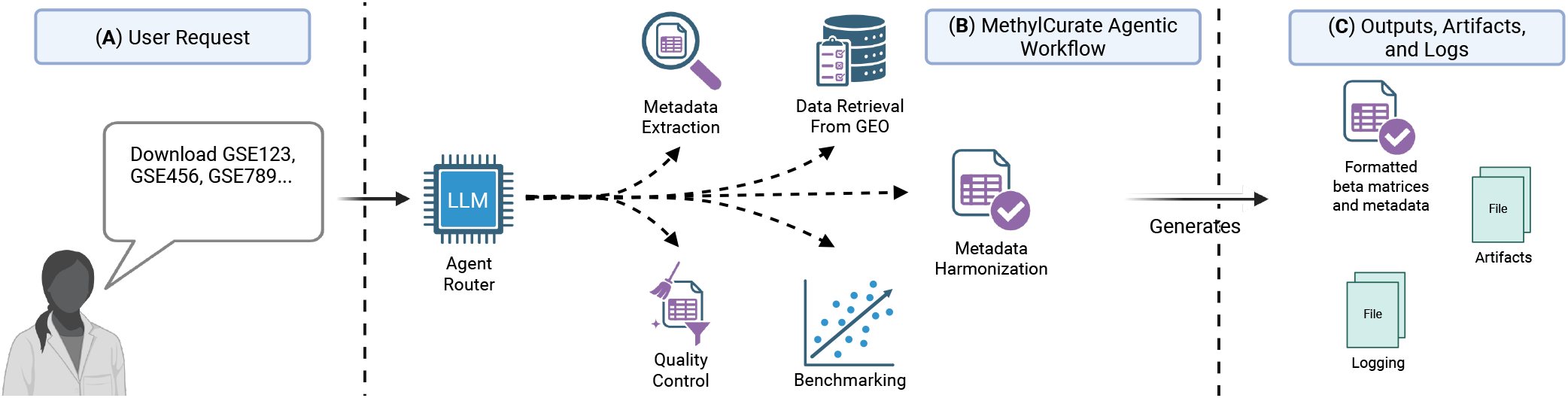

**Key Messages:**

- Automated curation of DNA methylation datasets from the Gene Expression Omnibus.
- Standardized preprocessing and metadata harmonization.
- Integrated benchmarking of epigenetic aging clocks.

## Introduction

Globally, human life expectancy has continued to rise, with a life expectancy at birth of 73.3 years in 2024 compared with roughly 64.9 two decades earlier (8). However, these gains in lifespan have not been matched by comparable increases in years lived free of disease and disability. Chronological age remains one of the strongest risk factors for chronic disease and mortality, yet it is an imperfect measure of an individual’s physiological state. Individuals of the same chronological age can differ substantially in health outcomes, motivating the use of biological age as a quantitative measure of underlying physiological condition.

DNA methylation (DNAm) is one of the most widely used modalities for developing models of biological age, commonly referred to as aging clocks. The most common type of aging clock is the “first-generation aging clock,” which is trained on cross-sectional data from healthy cohorts to predict chronological age. Although efforts are emerging to provide standardized datasets for model development and evaluation (3), prior work largely still depends on assembling DNAm data from public repositories such as NCBI Gene Expression Omnibus (GEO) (1), ArrayExpress (5), and EWAS Data Hub (9).

For researchers developing new clocks or evaluating existing ones, preparing DNAm data remains a major bottleneck. Public repositories often contain inconsistently structured metadata and study-specific supplementary files, all of which complicate automated dataset retrieval and harmonization and frequently require substantial manual effort. This heterogeneity also extends to processed DNAm matrices, which can differ across studies in preprocessing, formatting, and measurement representation. Recent resources such as BioLearn (10) and ComputAgeBench (3) address part of this problem, but important limitations remain. BioLearn supports a broader set of datasets, but metadata are not fully harmonized and not all studies are supported. ComputAgeBench is a static collection focused on blood DNAm data, which limits its utility for researchers interested in other tissues. Manual curation therefore remains common, despite being time-consuming, error-prone, and difficult to scale.

Here we present MethylCurate, a tool designed to support both curation of GEO DNAm datasets and evaluation of preexisting epigenetic aging clocks. MethylCurate streamlines the retrieval of DNAm data from GEO, harmonization of study metadata to relevant ontologies, construction of data into a unified format, and benchmarking of existing aging clocks without relying on labor-intensive manual extraction. By improving data accessibility, standardization, and reproducibility, MethylCurate enables researchers to gather, develop, and evaluate epigenetic aging clocks more efficiently.

### Implementation

MethylCurate is implemented in Python and exposed through a dialogue-driven interface. The framework combines a routing layer, deterministic processing modules, and LLM-assisted curation modules so that users can retrieve studies, generate standardized metadata, construct formatted beta matrices, and benchmark epigenetic aging clocks within a single workflow. Processing decisions, generated artifacts, and transformed outputs are logged throughout the workflow to preserve provenance and support reproducibility.

### Deterministic Modules

The deterministic components execute fixed computational procedures that do not require LLM-based reasoning. GEO retrieval is performed with the GEOparse Python library, which downloads study metadata, per-sample metadata, and per-sample DNAm data. In the event that per-sample DNAm data is missing, the interface can prompt the user to select the appropriate supplementary file containing the processed DNAm data.

Quality control is performed through a series of standard preprocessing steps. The workflow first determines whether methylation measurements are provided as beta values or M-values, converting them to beta values when necessary, and then applies sample- and CpG-level filtering. MethylCurate can subsequently evaluate epigenetic aging clocks through a wrapper around the PyAging library (2), enabling streamlined model evaluation across retrieved datasets.

### LLM-assisted Modules

Some curation tasks are difficult to solve reliably with rules alone, particularly when GEO metadata fields are inconsistently formatted or when supplementary files use study-specific layouts. For these tasks, MethylCurate uses LLM-assisted agents that generate structured outputs under explicit schema constraints.

#### Routing

Because the framework operates through a dialogue-driven interface, a routing component is used to determine which action should be performed in response to a user query. This router is implemented using LangGraph in combination with structured prompting and schema-constrained outputs. Based on the user’s request and the history of actions taken during the conversation, the router determines the most appropriate module to execute and can also recommend potential next steps.

#### Metadata Extraction

This agent infers how to extract sample-level metadata from GEO datasets when provided with a representative subset of metadata entries. To reduce hallucination during this process, we require structured outputs that specify the target fields along with regular expressions used for extraction. As a result, any extracted data must originate directly from the dataset. The extraction process is iterative: the proposed extraction rules are first applied to the full dataset, and the resulting output is evaluated for missing values or inconsistencies. If issues are detected, these failures are reported back to the LLM, which is then prompted to refine its extraction strategy.

#### DNA Methylation Data Extraction

GEO supplementary files often store processed DNAm data in study-specific formats, making purely rule-based parsing unreliable. MethylCurate therefore uses an iterative LLM-assisted procedure to parse candidate supplementary files, infer their formatting scheme, construct standardized beta matrices, and map samples back to the corresponding metadata. To do this, the agent examines random subsets of columns to infer column roles and identify layouts containing either beta values alone or paired beta-value and detection-value columns. The proposed parsing strategy is then applied to the full file and automatically evaluated for structural consistency, extraction completeness, and alignment between methylation data and sample metadata. When issues are detected, feedback on extraction errors is returned to the agent to support refinement.

#### Metadata Harmonization

Metadata fields such as sex, disease status, cell type, and tissue type often appear in heterogenous forms across studies. To harmonize these fields, MethylCurate first infers standardized human-readable labels using both the observed values and the dataset description as context. For disease, cell type, and tissue annotations, the inferred labels are used to query relevant biomedical ontologies, such as MONDO, CT, or Uberon respectively. The top k ontology matches are retrieved, and the LLM selects the most appropriate mapping while preserving the original study labels for provenance.

## Results

To assess the practical utility of our tool, we applied MethylCurate to 16 blood and brain-related GEO DNAm datasets (Fig. 1A; Supplementary Table 1). Supplementary materials summarize the vignette computing environment, dataset characteristics, file-selection decisions, quality control thresholds, metadata harmonization outputs, and interface screenshots (Supplementary Tables 1–4 and Supplementary Figures 1–4). MethylCurate successfully retrieved both study metadata and processed DNAm data for all 16 datasets. In contrast, BioLearn returned usable data for 4 of the 16 datasets.

**Figure 1.**
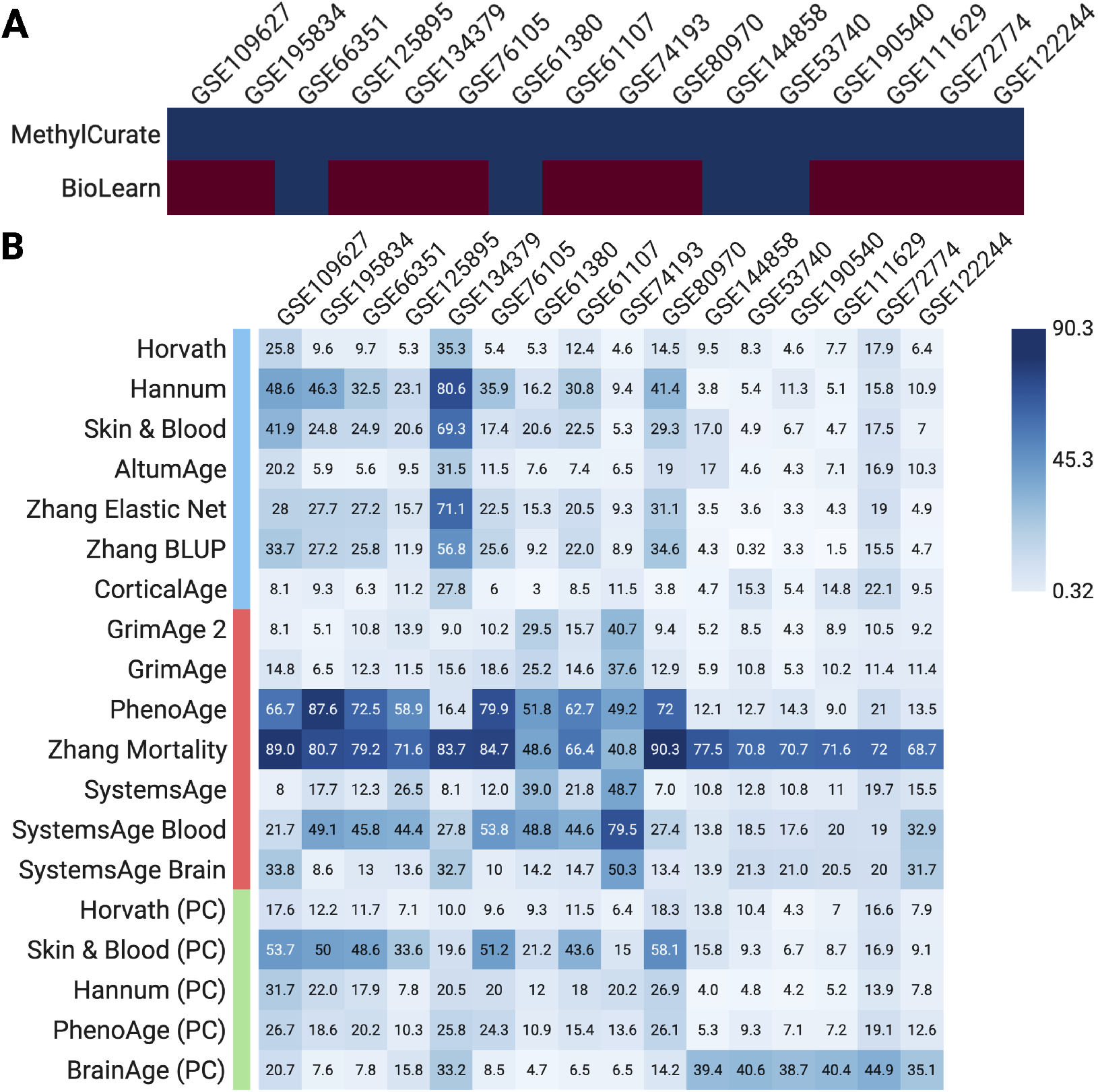
(A) Comparison of dataset retrieval success across 16 GEO datasets between MethylCurate and BioLearn. MethylCurate successfully retrieved and processed all evaluated datasets, whereas BioLearn supported a limited subset. Dataset accession IDs are shown on the x-axis. Red indicates failure to retrieve both metadata and methylation data, and blue indicates successful retrieval of both (B) Heatmap summarizing the performance of 19 epigenetic aging clocks across curated datasets. Values represent mean absolute error for each clock-dataset pair. Rows correspond to aging clock models and columns correspond to datasets. Both panels were generated using BioRender. The rows are color-coded to denote the type of model. Blue refers to the first-generation aging clocks (Horvath, Hannum, Skin&Blood, AltumAge, Zhang Elastic Net, Zhang BLUP, and CorticalAge), red refers to second-generation aging clocks (GrimAge 2, GrimAge, PhenoAge, ZhangMortality, SystemsAge, SystemsAge Blood, and SystemsAge Brain), and green refers to principal-component based aging clocks (Horvath (PC), Skin&Blood (PC), Hannum (PC), PhenoAge (PC), and BrainAge (PC). Both of these figures were generated using Biorender.

We next used the curated datasets to benchmark 19 epigenetic aging clocks through the unified evaluation workflow (Fig. 1B). Mean absolute error varied widely across model-dataset pairs, ranging from 0.32 to 90.3 years, indicating substantial heterogeneity in clock performance across cohorts. Rather than identifying a single universally best model, this analysis illustrates the importance of consistent multi-dataset benchmarking and shows that MethylCurate can move directly from data retrieval and metadata harmonization to comparative clock evaluation.

## Discussion

Public repositories contain a large number of potentially valuable DNAm datasets, but heterogeneity in metadata structure, sample annotation, and supplementary-file formatting makes these resources difficult to use at scale. MethylCurate addresses this bottleneck by transforming heterogeneous GEO studies into standardized inputs for reproducible benchmarking. Although existing tools for aging clock evaluation, including MethylCIPHER (7), BioAge (4), methylclock (6), ComputAgeBench (3), and BioLearn (10), provide useful benchmarking or biological-age analysis capabilities, MethylCurate complements these efforts by focusing on dataset retrieval, metadata harmonization, quality control, and integration with PyAging (2).

Several limitations remain. First, performance depends partly on GEO download speed and NCBI rate limits, both of which can lead to slow large-scale retrieval. Second, MethylCurate can occasionally confuse tissue and cell-type annotations when harmonizing GEO metadata (see Supplementary Table 4). This limitation could be addressed through improved prompting and ontology-mapping strategies, and it does not affect downstream clock performance because these labels are not used during clock evaluation. Third, MethylCurate currently focuses on GEO. Although GEO contains many of the DNAm datasets used in epigenetic aging research, extending support to repositories such as ArrayExpress and EWAS Data Hub would broaden the tool’s applicability.

Overall, MethylCurate provides a practical framework for converting heterogeneous DNAm studies into standardized datasets for epigenetic aging research. The dialogue-driven interface improves accessibility for users with varying levels of computational expertise, and the open-source design creates opportunities for continued improvement in prompting, extraction strategies, and repository support. MethylCurate may serve as a pivotal tool to accelerate the evaluation and development of epigenetic aging clocks.

## Data availability

All datasets analyzed in this study are publicly available from NCBI GEO. Source code, documentation, and example workflows are available at https://github.com/Travyse/methylcurate

## Author contributions statement

T.A.E. conceived the experiment, conducted the experiment, developed the tool, analyzed the results, and wrote the manuscript. T.A.E, L.S., and Q.L reviewed the manuscript.

## Acknowledgments

The authors thank the anonymous reviewers for their valuable suggestions.

