## Supplementary Information for "MethylCurate: Tool for Dataset Curation and Epigenetic Aging Clock Evaluation"

May 2026

#### Supplementary Methods

##### Computing Environment

The vignette described in the primary manuscript was run on a 2023 Apple M2 MacBook Pro with 32 GB of memory. MethylCurate was executed in a Docker container following the instructions provided in the project GitHub repository. The container was allocated 12 GB of memory through Docker Desktop. Large language model-assisted modules used Qwen3.5 397B through Ollama Cloud. No fine-tuning or domain-specific retraining was performed.

#### Supplementary Tables

Supplementary Table 1: Characteristics of DNA methylation datasets used in the vignette

| Accession Code | Sample Count | Age Range | Sex | Condition | Platform |
|---|---|---|---|---|---|
| GSE53740 | 384 | 34.00 - 93.00 | M (33.85%), F (40.36%) | Dementia | Illumina HumanMethylation450 |
| GSE61107 | 48 | 20.00 - 94.00 | M (70.83%), F (25.00%) | Dementia | Illumina HumanMethylation450 |
| GSE61380 | 33 | 21.00 - 73.00 | M (84.85%), F (15.15%) | Psychiatric | Illumina HumanMethylation450 |
| GSE66351 | 190 | 18.00 - 97.00 | M (46.84%), F (53.16%) | Dementia | Illumina HumanMethylation450 |
| GSE72774 | 508 | 35.10 - 91.90 | M (55.31%), F (44.69%) | Dementia | Illumina HumanMethylation450 |
| GSE74193 | 675 | -0.50 - 96.98 | M (63.85%), F (36.15%) | Psychiatric | Illumina HumanMethylation450 |
| GSE76105 | 68 | 66.00 - 94.00 | M (48.53%), F (51.47%) | Dementia | Illumina HumanMethylation450 |
| GSE80970 | 286 | 70.00 - 108.00 | M (37.76%), F (62.24%) | Dementia | Illumina HumanMethylation450 |
| GSE109627 | 162 | 70.00 - 95.00 | M (48.15%), F (51.85%) | Dementia | Illumina HumanMethylation450 |
| GSE111629 | 572 | 35.00 - 92.00 | M (56.47%), F (43.53%) | Dementia | Illumina HumanMethylation450 |
| GSE122244 | 70 | 45.00 - 82.00 | M (60.00%), F (40.00%) | Dementia | Illumina HumanMethylation850 |
| GSE125895 | 269 | 51.83 - 92.29 | M (55.39%), F (44.61%) | Dementia | Illumina HumanMethylation450 |
| GSE134379 | 808 | 54.00 - 103.00 | M (50.50%), F (49.50%) | Dementia | Illumina HumanMethylation450 |
| GSE144858 | 300 | 52.00 - 90.00 | M (40.00%), F (60.00%) | Dementia | Illumina HumanMethylation450 |
| GSE190540 | 90 | 60.34 - 83.63 | M (51.11%), F (46.67%) | Dementia | Illumina HumanMethylation850 |
| GSE195834 | 78 | 59.00 - 87.00 | M (52.56%), F (47.44%) | Dementia | Illumina HumanMethylationEPIC |

Supplementary Table 1 summarizes the 16 blood and brain DNAm datasets used in the vignette. For each dataset, the table reports accession code, sample count, age range, sex distribution, disease or condition, and methylation array platform. Sex percentages may sum to less than 100% where sex annotations are missing for some samples.

Supplementary Table 2: User-selected supplementary files for DNA methylation beta-matrix extraction

| Accession Code | Decision |
| --- | --- |
| GSE109627 | GSE109627_MTG_processed_GA_illumina_methylation.txt.gz |
| GSE111629 | GSE111629_PEGblood_450kMethylationDataBackgroundNormalized.txt.gz |
| GSE122244 | GSE122244_beta_LB.txt.gz<br>GSE122244_beta_LT.txt.gz<br>GSE122244_beta_M.txt.gz<br>GSE122244_beta_N.txt.gz<br>GSE122244_whole_blood.txt.gz |
| GSE125895 | GSE125895_matrix_processed.csv.gz |
| GSE134379 | GSE134379_processedSamples_cblAD.txt.gz<br>GSE134379_processedSamples_cblND.txt.gz<br>GSE134379_processedSamples_mtgAD.txt.gz<br>GSE134379_processedSamples_mtgND.txt.gz |
| GSE195834 | GSE195834_2-PD-processed.csv.gz |
| GSE72774 | GSE72774_datBetaNormalized.csv.gz |
| GSE74193 | GSE74193_GEO_procData.csv.gz |
| GSE66351 | — |
| GSE76105 | — |
| GSE61380 | — |
| GSE61107 | — |
| GSE80970 | — |
| GSE144858 | — |
| GSE53740 | — |
| GSE190540 | — |

Supplementary Table 2 lists the user-selected supplementary files used when processed per-sample DNAm data were not directly available from GEO sample records. In these cases, MethylCurate prompted the user to identify the file most likely to contain the processed DNAm beta matrix. A dash indicates that no user-selected supplementary file was required.

Supplementary Table 3: Quality control thresholds used in the vignette

| Parameter | Meaning | Value |
| --- | --- | --- |
| dnam_cutoff | Threshold used to remove samples that do not have a maximum DNAm above this value. | 0.96 |
| sample_level_missing_cutoff | Maximum allowed fraction of missing values per sample. | 0.1 |
| cpg_level_missing_cutoff | Maximum allowed fraction of missing values per CpG site. | 0.2 |
| correlation_cutoff | Minimum allowed mean inter-array correlation per sample. | 0.9 |

Supplementary Table 3 reports the quality control thresholds used in the vignette. These values correspond to the default MethylCurate settings and were applied across retrieved datasets before downstream clock evaluation.

Supplementary Table 4: Harmonization of Metadata Across 16 Datasets

| Concept | Grouping | Harmonized Value | Raw Value |
|---|---|---|---|
| Disease | Neurodegenerative Disease | Alzheimer Disease | AD, Alzheimers Disease, Alzheimer's disease, Alzheimer |
| | | Corticobasal Degeneration Disorder | CBD |
| | | Frontotemporal Dementia | FTD |
| | | Frontotemporal Dementia with Motor Neuron Disease | FTD/MND |
| | | Progressive Supranuclear Palsy | PSP |
| | | Parkinson Disease | PD, Parkinson's disease (PD), Parkinson's disease |
| | Psychiatric Disorder | Schizophrenia | 2, schizophrenia, Schizo |
| | Cognitive Disorder | Cognitive Impairment | mild cognitive impairment, Case |
| | Unknown | Unknown | Unknown |
| | Control | Control | 1, Control, CTRL, ND |
| Tissue | Leukocyte | Peripheral Blood Mononuclear Cell | Peripheral Blood, peripheral blood leukocytes |
| | | B Cell | lymphocytes B |
| | | T Cell | lymphocytes T |
| | | Monocyte | monocytes |
| | | Neutrophils | neutrophils |
| | Blood | blood | Whole Blood |
| | Brain | Frontal Cortex | Frontal Cortex |
| | | Prefrontal Cortex | Brain Frontal Cortex |
| | | Occipital Cortex | occipital cortex |
| | | Temporal cortex | temporal cortex |
| | | Dorsolateral Prefrontal Cortex | DLPFC |
| | | Superior Temporal Gyrus | Superior Temporal Gyrus |
| | | Middle Temporal Gyrus | brain; middle temporal gyrus, MTG |
| | | Entorhinal Cortex | ERC |
| | | Cerebellum | CRB, CBL |
| | | Hippocampal Formation | HIPPO |
| | - | **Normal Brain tissue** | Normal Brain tissue |
| | - | **Parkinson's disease Affected Brain Tissue** | Parkinson's disease Affected Brain Tissue |
| Cell Type | - | Glial Cell | glial cell |
| | - | Neuron | neuron |
| | - | Bulk | bulk |
| | - | B cell | B cell |
| | - | T cell | T cell |
| | - | monocyte | monocyte |
| | - | neutrophil | neutrophil |
| | - | blood cell | blood |
| Sex | - | Female | FEMALE, F |
| | - | Male | MALE, M, MM |
| | - | Unknown/Other | UNKNOWN, ?, Unsure |

Supplementary Table 4 reports the metadata harmonization output across the 16 curated datasets. The "Concept" column denotes the metadata field being harmonized, the "Grouping" column provides a broader category used to organize harmonized labels, the "Harmonized Value" column reports the selected standardized label or ontology-aligned term, and the "Raw Value" column lists the original dataset-specific annotations mapped to that value. When an ontology-derived term could not be found by MethylCurate, an LLM-inferred best-guess label was used. The bold values are the LLM-inferred best guesses.

### Supplementary Figures

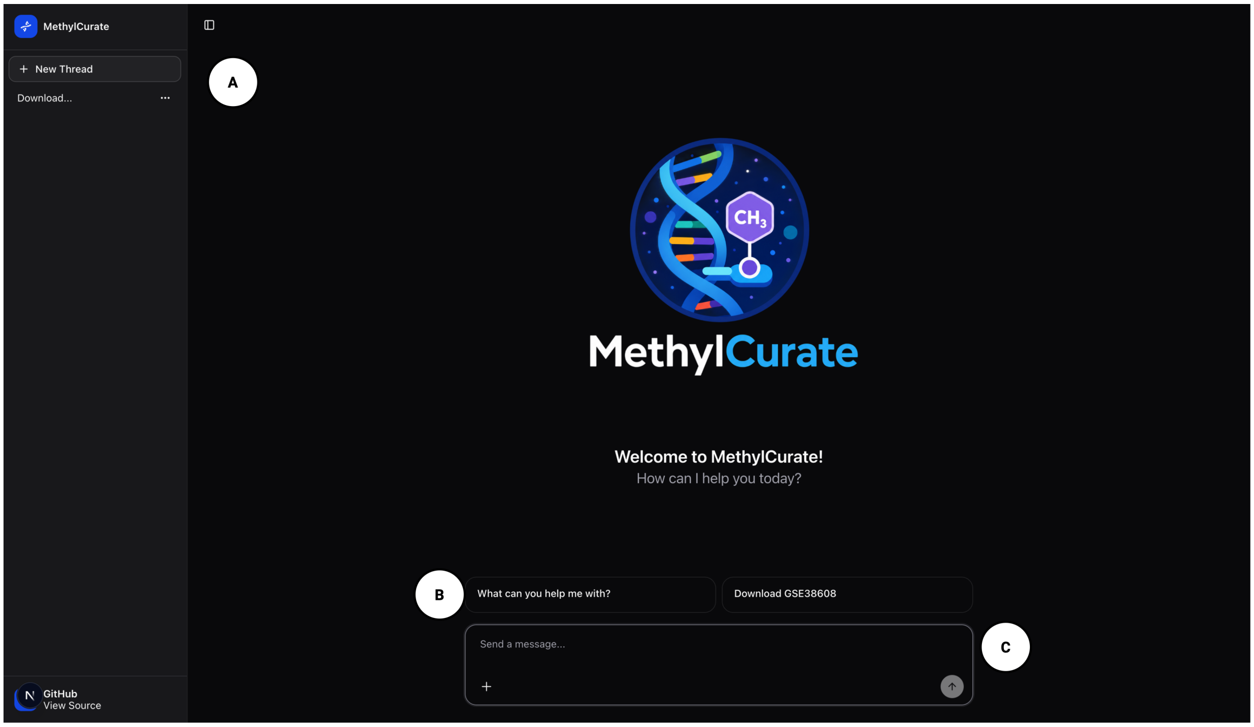


Supplementary Figure 1: **MethylCurate dialogue-driven user interface.** (A) The sidebar displays existing conversation threads and includes a "New Thread" button for starting a new session. (B) The central panel provides predefined prompt options that users can select to initiate common workflows. (C) The chat box allows users to submit free-text queries.

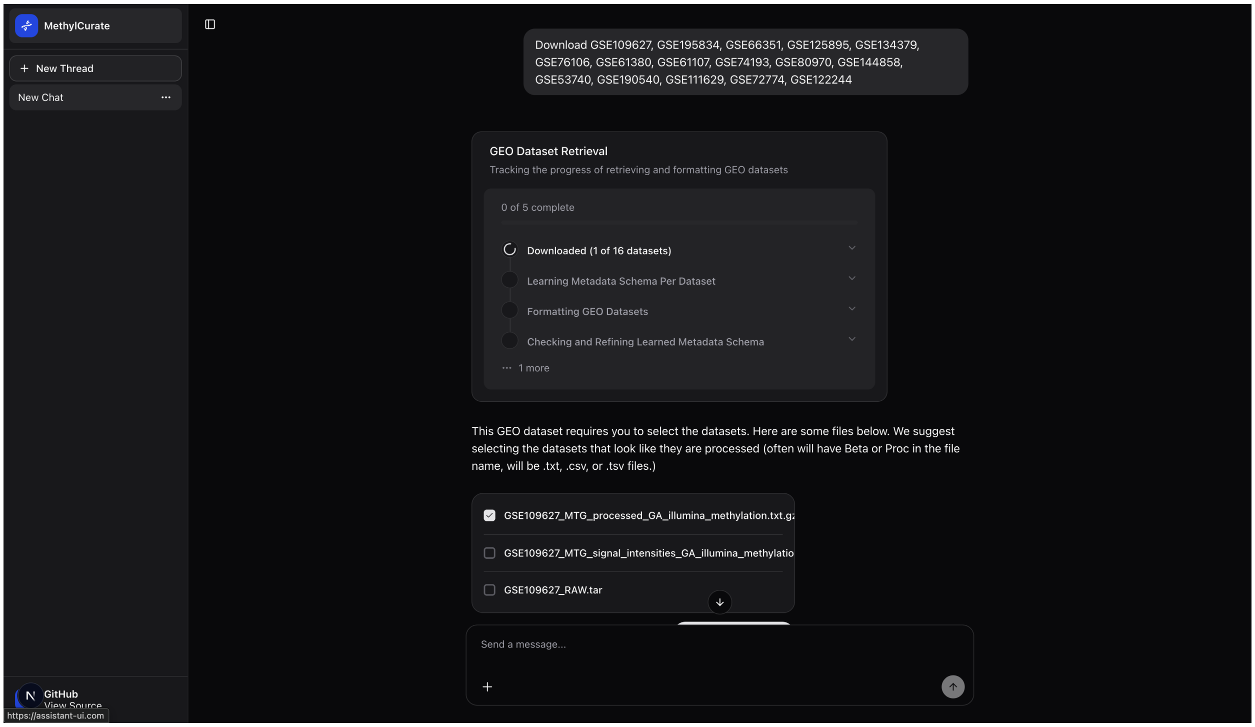


Supplementary Figure 2: **Human-in-the-loop supplementary file selection during GEO dataset retrieval.** When per-sample DNAm data are not directly available, MethylCurate prompts the user to select the supplementary file most likely to contain the processed beta matrix.

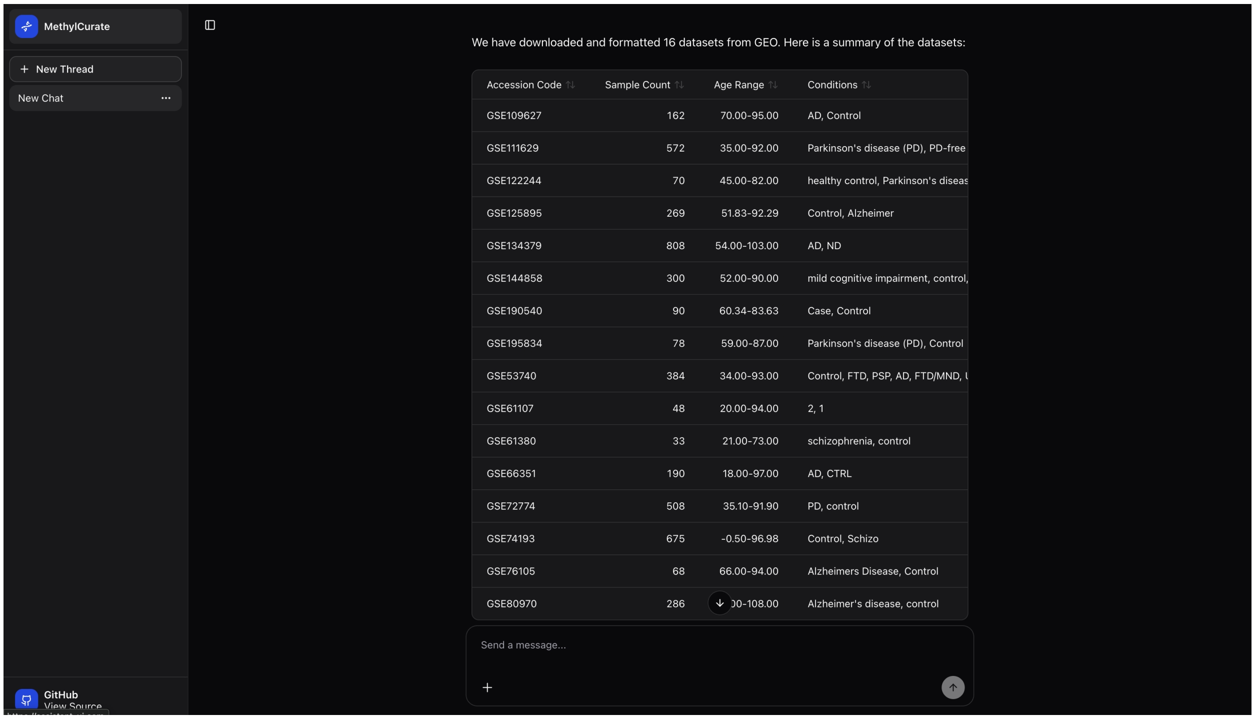


Supplementary Figure 3: **Dataset retrieval summary after GEO processing.** After dataset retrieval, MethylCurate displays summary statistics for each successfully retrieved dataset, including accession code, sample count, age range, and condition labels.

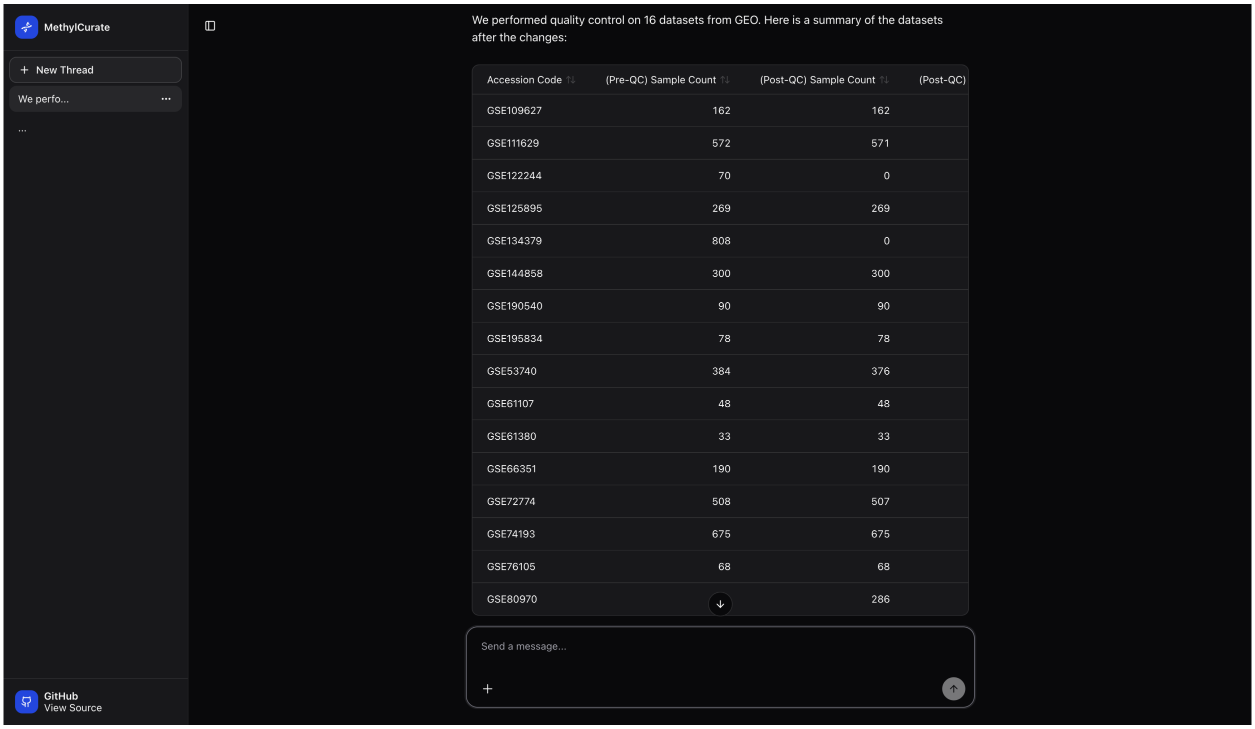


Supplementary Figure 4: **Quality control summary after dataset filtering.** After quality control, MethylCurate displays pre-QC and post-QC sample counts, along with post-QC CpG feature counts for each dataset.
